# Clemastine induces an impairment in developmental myelination

**DOI:** 10.1101/2021.12.14.472570

**Authors:** Ana Palma, Juan Carlos Chara, Amaia Otxoa-Amezaga, Anna Planas, Carlos Matute, Alberto Pérez-Samartín, María Domercq

## Abstract

Abnormalities in myelination are associated to behavioral and cognitive dysfunction in neurodevelopmental psychiatric disorders. Thus, therapies to promote or accelerate myelination could potentially ameliorate symptoms in autism. Clemastine, a histamine H1 antagonist with anticholinergic properties against muscarinic M1 receptor, is the most promising drug with promyelinating properties (Mei et al., 2014). Clemastine penetrates the blood brain barrier efficiently and promotes remyelination in different animal models of neurodegeneration including multiple sclerosis, ischemia and Alzheimer’s disease. However, its role in myelination during development is unknown. We showed that clemastine treatment during development increase oligodendrocyte differentiation in both white and gray matter. However, despite the increase in the number of oligodendrocytes, conduction velocity of myelinated fibers of *corpus callosum* decreased in clemastine treated mice. Confocal and electron microscopy showed a reduction in the number of myelinated axons and nodes of Ranvier and a reduction of myelin thickness in *corpus callosum*. To understand the mechanisms leading to myelin formation impairment in the presence of an excess of myelinating oligodendrocytes, we focused on microglial cells which also express muscarinic M1 receptors. Importantly, the population of CD11c^+^microglia cells, necessary for myelination, as well as the levels of insulin growth factor-1 decrease in clemastine-treated mice. Altogether, these data suggest that clemastine impact on myelin development is more complex than previously thought and could be dependent on microglia-oligodendrocyte crosstalk. Further studies are needed to clarify the role of microglia cells on developmental myelination.

## 1. INTRODUCTION

Myelin directs node of Ranvier formation, enable rapid saltatory conduction, communicates with axons via the periaxonal space and provides metabolic support to them. Myelin, as other structures in brain, is subject to plastic changes in response to electrical activity that allow to fine tune circuits involved in motor learning and memory (McKenzie et al., 2014). Changes in the number or structure of myelin sheaths, node of Ranvier and periaxonal space modulate axon conduction velocity. Myelin formation includes diverse and very controlled events: migration and proliferation of oligodendrocyte precursor cells (OPCs), recognition of target axons, differentiation of OPCs into mature oligodendrocytes, membrane outgrowth, axonal wrapping, and myelin compaction. Myelination takes place during the developmental critical period of plasticity, when experience can rapidly change cortical structure and function. Morever, early GABAergic communication between OPCs and interneurons determines interneuron maturation and myelination (Fang et al., 2021). Thus, myelination could have a more active role in circuit refinement and maturation during development. The importance of myelin for circuit function is made apparent by the severity of neurological diseases associated with its disruption. Abnormalities in myelination during development causes cognitive dysfunction and are associated to behavioral disturbances in neurodevelopmental psychiatric disorders, as reported in several animal models of syndromic Autism Spectrum Disorders (ASD). For instance, the normal process of myelination is delayed in Contactin-associated protein-like 2 (Caspr2) knock out mice (Scott et al., 2019); and a mouse model of Pitt–Hopkins syndrome (PTHS), a syndromic form of ASD, showed reductions in mature oligodendrocyte numbers and myelination (Phan et al., 2020). However, to date, there is no therapy to compensate or alleviate neurodevelopmental abnormalities or delays of myelination. The fact that myelin is transiently altered in these neurodevelopmental disorders point to pro-myelinating drugs as a promising therapeutic approach.

Clemastine is to date the most promising pro-myelinating drug. Clemastine is an anti-histaminic compound with anti-muscarinic properties against the muscarinic receptor 1 (M1R) (de Angelis et al., 2012). Previous reports confirm that muscarinic M1 receptors are expressed by oligodendroglial cells and may modulate OPC proliferation and differentiation (de Angelis et al., 2012; Deshmukh et al., 2013; Mei et al., 2014). Morevoer, M1Rs antagonist clemastine, as well as other antagonists such as benztropine, promote remyelination in animal models of multiple sclerosis (Deshmukh et al., 2013; Mei et al., 2014), after hypoxia (Cree et al., 2018; Wang et al., 2018) and prevent demyelination secondary to aging and in Alzheimer’s disease (Wan et al., 2020; Chen et al., 2021). However, little is known about the effect of clemastine in the absence of myelin abnormalities or demyelination. Here, we analyzed the impact of clemastine in myelination during development. Surprisingly, clemastine decreased axon conduction velocity despite the increase in oligodendrogenesis. Our data point to microglia as another target of clemastine involved in developmental myelination.

## MATERIALS AND METHODS

### Animal treatments

Experiments were performed in C57BL/6 wild-type mice and CD11c^+^ eYFP mice (generously provided by the Laboratory of Cerebrovascular Research, IIBB-CSIC, Barcelona). All experiments were performed according to the procedures approved by the Ethics Committee of the University of the Basque Country (UPV/EHU). Animals were handled in accordance with the European Communities Council Directive. Animals were kept under conventional housing conditions (22 ± 2°C, 55 ± 10% humidity, 12-hour day/night cycle and with *ad libitum* access to food and water) at the University of the Basque Country animal unit. All possible efforts were made to minimize animal suffering and the number of animals used. C57BL/6 wild-type mice were daily treated with vehicle or clemastine (#1453 Tocris; 50mg/kg solved in 10% DMSO) by intraperitoneal injections from postnatal day 5 (P5) to P21. CD11c^+^ eYFP mice were treated from P5 to P10 following the same protocol.

### Immunohistochemistry

Mice were transcardially perfused with 4% p-formaldehyde in 0.1 M sodium phosphate buffer (PBS) (pH 7.4) and post-fixated in 4% PFA for 4 hours. Tissue was cut using a Microm HM650V vibratome (40 μm). After blocking with 4% normal goat serum in PBS plus 0.1% Triton X-100 (blocking solution) for 1h at room temperature, free-floating sections were incubated with primary antibodies overnight at 4°C in blocking solution. The following antibodies were used: anti-APC (1: 500 #OP80 Sigma), anti-Olig2 (1:1000 #MABN50 Sigma), anti-Caspr (1:500 #MABN69 Sigma), anti-MBP (1:1000 #808401 Biolegend), anti-NaV1.6 (1:500 #ASC-009 Alomone), anti-iNOS (1:500 #610329 BDBiosciences), anti-PDGFRα (1:250 #sc-398206 SantaCruz Biotechnology), anti-Iba1 (1:500 #019-19741 Wako). Tissue sections were washed and incubated with the appropriate secondary antibodies: AlexaFluor-488, −594, and −647 anti-rat, anti-rabbit, anti-guinea pig, and anti-mouse IgG secondary antibodies (Invitrogen) for 1h at room temperature. Cell nuclei were stained using DAPI (1: 1000 Vector Laboratories).

Images were acquired with the same settings for all samples within one experimental group. Zeiss LSM800, Zeiss LSM880 Airyscan and Leica TCS SP8 confocal microscope were used to acquire the images. To quantify the differentiation of oligodendrocytes, z-stack images (40X) were taken using a Leica TCS SP8 confocal microscope. Results were expressed as a percentage of APC^+^ and PDGFR^+^ *vs* the total number of olig2^+^ oligodendrocytes. To quantify the number and length of nodes of Ranvier and paranodes in *corpus callosum* and cerebral cortex, high magnification (63X) z-stack images were taken using a Zeiss LSM800 confocal microscope and analysis was performed in maximal projections obtained with ImageJ software (NIH). Node and paranodes length were only measured when a node and its flanking paranodes were completely defined using the antibodies to Caspr and NaV1.6. To measure internode length in cerebral cortex, high magnification (40X) z-stack images were taken using a Zeiss LSM800 confocal microscope. Individual internodes were traced as MBP^+^ processes flanked by Caspr^+^ paranodes and the lenght analyzed using ImageJ software. To quantify microglia Iba1, CD68 and iNOS expression, high magnification (63X) images were taken using a Leica TCS SP8 super resolution microscope. Immunoreactivity of Iba1 and iNOS was determined by applying default threshold with the ImageJ software and was normalized to area. CD68^+^ puncta were quantified and normalized to the number of cells. Analysis was performed in at least three different sections of n = 3 mice. To analyze the morphology of Iba1^+^ microglia from cerebral cortex, high magnification (63X) z-stack images were taken with Zeiss LSM800 confocal microscope. Individual microglia cell was firstly skeletonized to perform *Sholl* analysis (ImageJ plugin) as well as branches and junctions quantifications (at least 20-25 cells, n= 6 animals from two different experiments). In all cases, data come from n=6 animals from two different litters.

### Electron Microscopy

Mice were perfused with 4% p-formaldehyde, 2.5% glutaraldehyde and 0.5% NaCl in phosphate buffer, pH7.4. The brains were postfixed with the same fixative solution overnight at 4°C. The tissue was sagittally cut using a Leica VT 1200S vibrating blade microtome (Leica microsystems) to obtain 200 μm-thick sections. Tissue sections were postfixed in 2% OsO4, dehydrated in ethanol and propylenoxide, and embedded in EPON (Serva) for 24h at 60°C. Ultrathin 50 nm sections were obtained using a Leica Ultracut S ultramicrotome (Leica) and contrasted with 4% uranyl acetate (30 min) and lead citrate (6 min). High resolution electron microscope images were taken using a Zeiss EM900 electron microscope (Zeiss). Sectioning, imaging and analysis were carried out by an experimenter blind to the treatment group. Image analysis was performed using ImageJ (NIH). The g-ratio of myelinated axons was calculated as the ratio of the inner to the outer radius of the myelin sheath using 8,000x magnitication images from at least 100-150 axons from 60 images per animal. The number of myelinated axons was quantified using 1,000x magnification images.

### Microglia culture

Primary mixed glial cultures were prepared from the cerebral cortex of neonatal rats and mice (P0–P2). After 10–15 days in culture, microglia were isolated by mechanical shaking (400 rpm, 1h) and cultured as previously described (Domercq et al., 2007).

### Cytosolic Ca2^+^ Imaging

To measure cytosolic [Ca^2+^], cells were loaded with Fluo-4 AM (1 mM; Molecular Probes, Invitrogen) in incubation buffer for 30 min at 37°C and washed (20 min). Images were acquired through a 63X objective by an inverted LCS SP8 confocal microscope (Leica, Germany) at an acquisition rate of 1 frame/15 s during 5 min. For data analysis, an homogeneous population of 15–25 cells was selected in the field of view and cell somata selected as ROIs. Background values were always subtracted and data was expressed as F/F0 ± SEM (%) in which F represents the fluorescence value for a given time point and F0 represents the mean of the resting fluorescence level.

### Microglia sorting and qPCR analysis

Total brain was homogenated and digested mechanically and enzymatically, and myelin debris was removed through a continuous 60X percoll gradient. Microglia was sorted using CD11b (#101205, 1:100; Biolegend) and CD45 (#103134, 1:100; Biolegend), to distinguish between resident microglia (CD11b^+^/CD45low) and macrophages (CD11b^+^/CD45high; Szulzewsky et al., 2015). Microglia was directly collected in RNA lysis buffer and total RNA was extracted using the RNeasy Plus Micro Kit (#74034) (Quiagen). RNA concentration and integrity were analyzed with the collaboration of the General Genomics Service Sequencing and Genotyping Unit from the UPV/EHU. A battery of microglia genes (see table 1) associated to microglia DAM signature, inflammatory reaction and/or phagocytosis was analysed. Gene expression was analysed using a 96.96 Dynamic Array™ integrated fluidic circuit (Fluidigm) real-time PCR and GenEx software. Results were depicted as relative gene expression according to the ∆∆Cq method (2−∆∆Ct) and expressed in base 2 logarithmic scale.

**Table 1.**
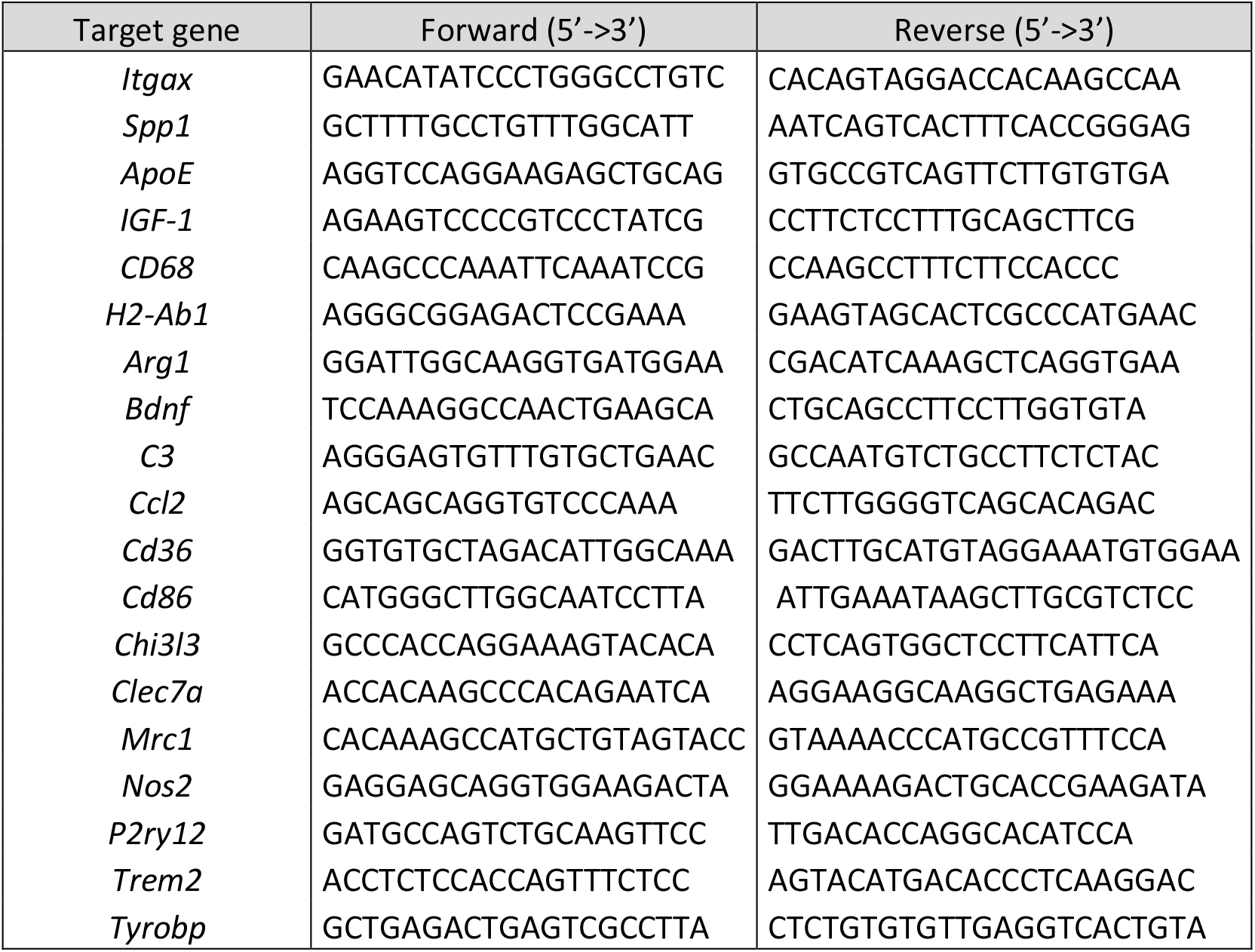
Sequences for mouse primers used for qPCR

### Electrophysiology

Mice were anesthetized and decapitated and the brain was rapidly removed and placed in ice-cold (4◦C) cutting artificial cerebrospinal fluid (ACSF) containing (in mM): 215 Sucrose, 2.5 KCl, 26 NaHCO3, 1.6 NaH2PO4, 20 Glucose, 1 CaCl2, 4 MGCl2, and 4 MgSO4 bubbled with a mixture of 95% O2/5% CO2. Coronal slices, 400-μm thick, were cut on a Leica VT1200S vibratome and transferred to a warmed (~36°C) solution of normal ACSF (nACSF) containing (in mM): 124 NaCl, 2.5 KCl, 10 Glucose, 25 NaHCO3, 1.25 NaH2PO4, 2.5 CaCl2, and 1.3 MgCl2 for recovery (45 min). Compound action potentials (CAPs) were evoked by electrical stimulation of the *corpus callosum* with a bipolar electrode (CE2C55, FHC, USA) and were recorded with a pulled borosilicate glass pipette (1.6 MΩ resistance) filled with NaCl 3 M within the contralateral *corpus callosum*. Stimulation intensities ranged from 30 to 3000 μA (100 μs pulses, Master-8, AMPI, Israel). Input-output curves were generated by recording the amplitudes of myelinated N1 and partially myelinated fibers N2 as a function of stimulation intensity. Conduction velocity values for N1 and N2 fibers were calculated as the slope of a straight line fitted through a plot of the distance between the recording and stimulating electrodes versus the response latency (time to N1 and N2 respectively). Recordings were perfomed at 4 different distances from the recording electrode (500, 1000, 1500 and 2000 μm). Three responses were averaged for each measurement. Peak amplitudes and latencies were calculated using custom written routines in pCLAMP 10.0 (Molecular Devices, USA).

### Statistical analysis

Data are presented as mean ± s.e.m. with sample size and number of repeats indicated in the figure legends. Statistical analysis was performed using GraphPad Prism statistical software (version 7.0; GraphPad software). Comparisons between two groups were analysed using paired Student’s two-tailed t-test. Statistical significance was considered at *p* < 0.05.

## 3. RESULTS

### 3.1 Clemastine increases the number of mature oligodendrocytes in developing brain

To determine whether clemastine treatment *in vivo* promotes myelination during development, we treated mice with clemastine (50mg/kg/day, i.p.) or vehicle from postnatal day 5 to 20, a time window coincident with myelination and within the critical period of brain plasticity (Fig. 1A). We first quantified the population of oligodendrocyte progenitor cells (OPCs) and mature oligodendrocytes (OLs) using antibodies against PDGFRa and CC1 (anti-APC), respectively. We performed blinded cell counts in three different areas, *corpus callosum* (CC), retrosplenial cortex (Rsp Ctx) and somatosensorial cortex (SSp Ctx), on anatomically equivalent brain sections and normalized our counts using the pan-OL marker Olig2. As showed in vitro (Mei et al., 2014, 2016), clemastine induced a significant increase in the proportion of APC^+^ OLs in *corpus callosum* as well as in the cerebral cortex (Fig. 1B). In parallel, we detected a decrease in the proportion of PDGFR^+^ OPCs in *corpus callosum*. In contrast, no change in the number of PDGFR^+^ OPCs was detected in cerebral cortex (Fig. 1C). These results indicate that the treatment with clemastine during development promotes the maturation of the oligodendroglial lineage by increasing the percentage of APC^+^ oligodendrocyte, particularly in the *corpus callosum*.

**Figure 1.**
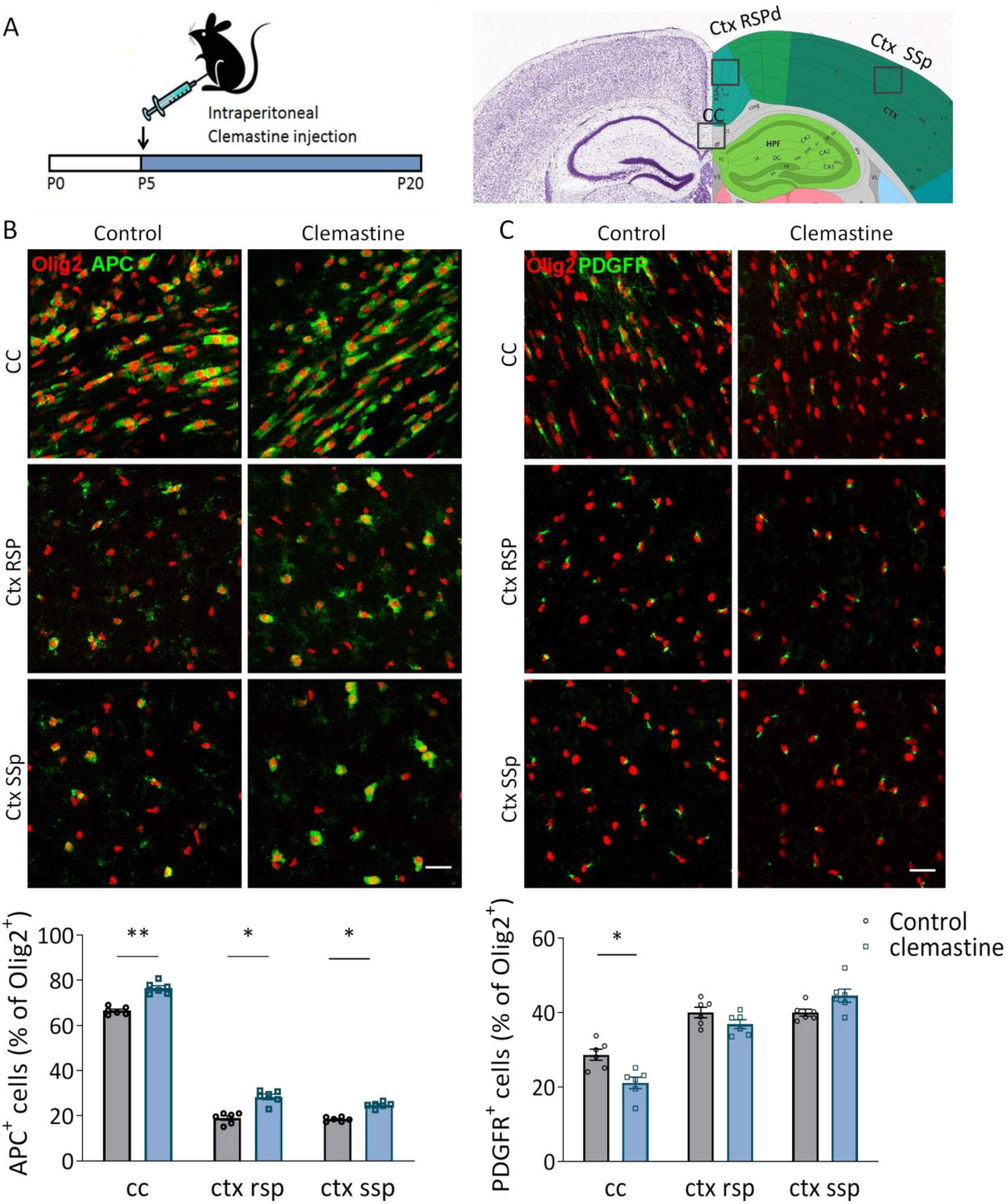
**A)** Schematic representation of clemastine treatment (50μg/kg; i.p.) during development. The analysis was performed in corpus callosum (cc), retrospinal cortex (Ctx rsp) and primary somatosensory cortex (ctx ss). **B, C)** Compressed confocal z stack images of APC^+^ mature oligodendrocytes (B; green) and PDGFR^+^ oligodendrocyte precursor cells (C; OPCs; green) in cc, ctx rsp and ctx ss of control and clemastine-treated mice. Below, average of the percentage of mature OLS (B) or OPCs (C) vs total Olig2^+^ oligodendroglial cells (Olig2^+^) (n= 6 mice per experimental group). Scale bar = 50 μm. *p<0,05, **p<0,01.

### 3.2 Conduction velocity of the interhemispheric fibers from *corpus callosum* decreased after clemastine treatment

As clemastine induced a clear increase in OL differentiaton in the *corpus callosum*, we hypothesized that this would translate into accelerated myelination and a subsequent increase in the conduction velocity in whiter matter. To test this idea, we measured the propagation of compound action potential (CAPs) in the *corpus callosum* by performing electrophysiological recordings in acute coronal brain slices. CAPs were evoked by a bipolar stimulating electrode and recordad by a field electrode placed at varying distantces across the *corpus callosum*. Surprisingly, our results revealed that action potentials transmission is significantly slower in myelinated axons (N1) of clemastine-treated mice compared to control mice (Fig. 2, p<0.01, n= 7-9). In contrast, the conduction velocity in unmyelinated axons (N2) is similar between groups (Fig.2, n= 7-9) Examining the relationship between stimulus intensity and response magnitude (input-output curve) revealed that myelinated axons in clemastine-treated mice tend to be more excitable than those in control mice (Fig. 2C). Therefore, we concluded that clemastine induced a delay in action potential propagation despite the effect in OL differentiation.

**Figure 2.**
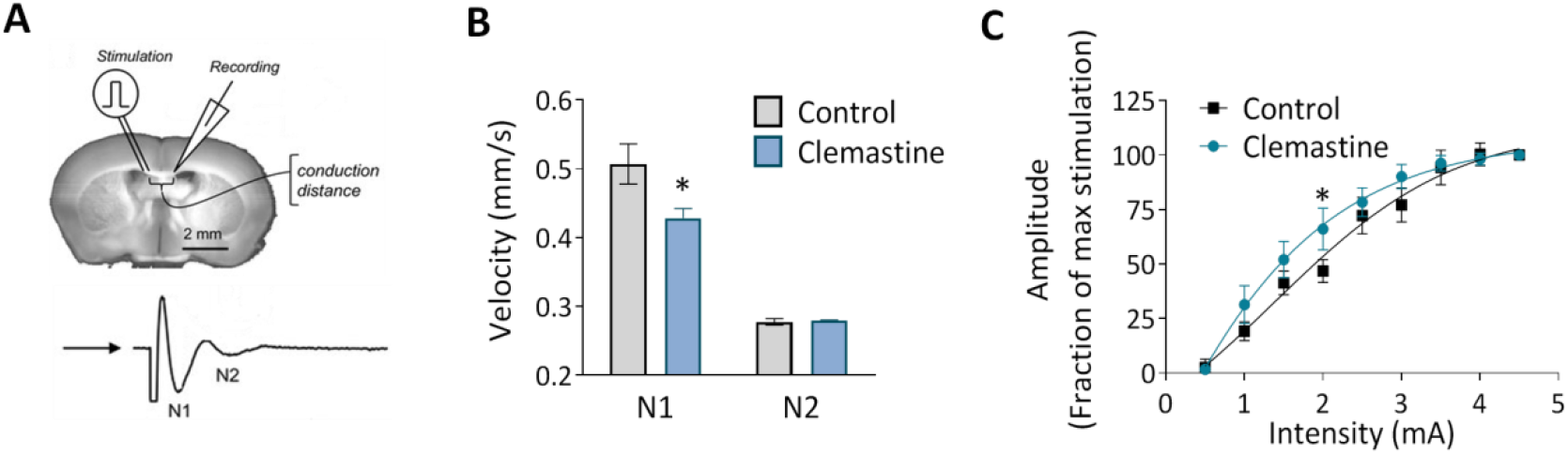
**A)** Schematic representation of compound action potentials (CAPs) recording in corpus callosum of 400μm coronal slices. **B)** Conduction velocity corresponding to N1 and N2 peaks in vehicle (n= 12) and clemastina-treated mice (n= 10). For each mice, the average measure come from at least four different sections recorded three times. **C)** The input-output relationship for N1 myelinated fibers is left-shifted in clemastine-treated corpus callosum (n= 10-12). *p<0,05

### 3.3 Clemastine administration during development reduces the number of nodes of Ranvier and myelinated axons in *corpus callosum*

Since the increase in the percentage of mature oligodendrocyte is not accompanied by an increase in the conduction velocity of N1 fibers in *corpus callosum*, myelination was further studied by performing histological analyses both by confocal and transmission electron microscopy. The speed of action potential conduction depends on the length and the number of the nodes of Ranvier (Arancibia-Cárcamo et al., 2017), in addition to myelin sheath number and structure, and node length is plastic and refined in response to electrical activity. We immunolabeled coronal brain sections with antibodies against contactin-associated protein (caspr) and sodium-gated channels 1.6 (Nav1.6) to visualize paranodes and nodes of Ranvier respectively. By identifying regions of dense Nav1.6 staining that were clearly flanked by abutting CASPR^+^ paranodes, we quantified the length of individual nodes within *corpus callosum* in control and clemastine-treated mice. We did not find any change in node length (Fig. 3A). However, we detected a significant reduction in the number of nodes of Ranvier in clemastine-treated mice (Fig. 3A). We also measured the length of internodes that were flanked on each end by contactin-associated protein (caspr^+^) paranodes. We found that clemastine treatment tend to increase internode length, although the differences were not significant (Fig. 3B).

**Figure 3.**
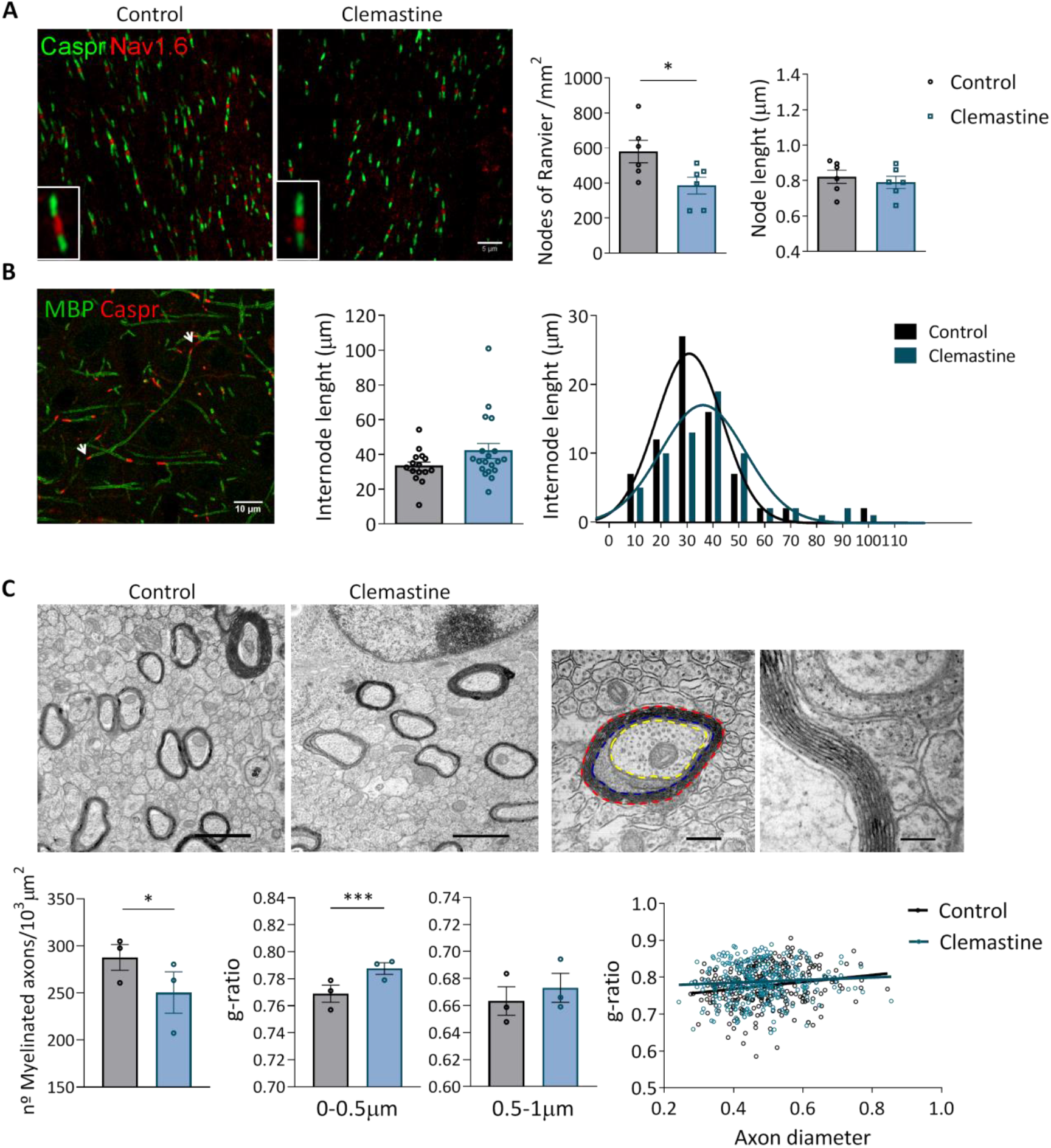
**A)** Representative confocal images of nodes of Ranvier (Nav1.6; red) and paranodes (Caspr; green) in corpus callosum of control and clemastine treated mice. Scale bar = 5 μm. Rigth, histograms shows average node of Ranvier number and length in control and clemastine-treated mice (n= 6 per experimental group). **B)** Representative confocal images of MBP^+^ myelin internodes (green) flanked by two Caspr^+^ paranodes (red; arrows). *Right*, average and cumulative internode length distribution in control and clemastine-treated mice (n= 6 per group). **C)** Representative electron microscopy images of myelinated axons within corpus callosum in control and clemastine treated mice. Graphs represent the average of myelinated axons (at least 60 different images per animal from n= 3 mice per group), the mean g-ratios for small and bigger axons (100-150 axons in control and clemastine-treated mice) and the distribution of g-ratios for control and clemastine-treated mice (n=3 mice per group).

The high density of myelin sheaths in corpus callosum made it difficult to look for differences using confocal microscopy. So, we next used transmission electron microscopy (TEM) to visualize myelination. TEM images were taken of the *corpus callosum* directly above the dorsal hippocampus from anatomically equivalent tissue sections from control and clemastine-treated mice. We observed that the number of myelinated axons, quantified by TEM, was reduced after clemastine treatment (Fig. 3C).

Myelin thickness was assessed by g-ratio calculation (axon diameter/total outer diameter of myelinated fiber). G-ratio analysis reveals thinner myelin sheaths in small caliber axons (0-0.5 μm) of treated animals, while no differences were found in axons from 0.5 to 1 μm diameter, indicating that the changes observed in myelin thickness depends on axons diameter (Fig. 3C).

Taken together, our findings suggest that, although clemastine treatment during development initially enhances oligodendrocyte differentiation, functional myelination is impaired or delayed.

### 3.4 Clemastine treatment modifies the expression activation markers and morphology of microglia in developing brain

In addition to electrical activity, a new player in developmental and adaptive myelination is microglial cells (Geraghty et al., 2019; Wlodarczyk et al., 2017). *Chmr1* gene, the receptor mediating clemastine-effect in OLs, is also expressed in microglia, as reported from previous RNAseq analysis (Fig. 4A; Zhang et al., 2014). Indeed, cultured microglia cells exposed to muscarine showed a significant increase in cytosolic calcium, as revealed with Fluo4-calcium imaging (Fig. 4B). We hypothesized that the impairment in myelination during development could be related to the interaction of clemastine with microglial cells. So, we next analyzed by immunohistochemistry microglia morphology and activation markers.

**Figure 4.**
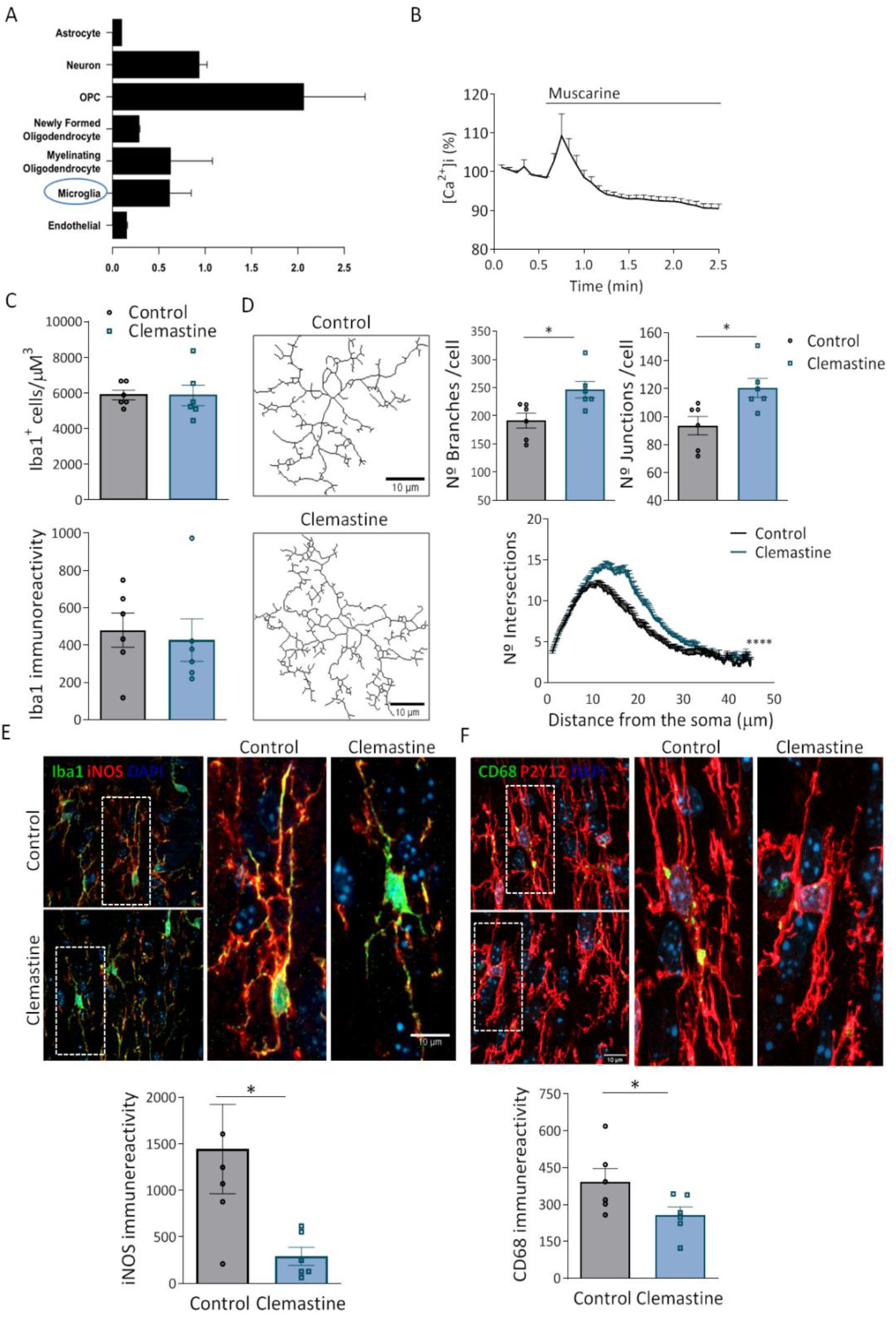
**A)** Graph showing *Chmr1* gen expression in all CNS cell types. Note that *Chmr1* is also expressed in microglia. Data was taken from BrainRNA-seq database (Zhang et al. 2014). **B**) Recordings of Ca^2+^ responses in microglia *in vitro* following application of muscarine (100 μM; arrow) using Fluo-4. Data represent average ± SEM of values obtained from more than 50 cells from 3 different cultures. **C)** Quantification of the number of Iba1+ microglia cells as well as Iba1 staining intensity in corpus callosum of control and clemastine-treated mice. **D)** Representative skeletonized images of microglia used for morphological analysis. Scale bar = 10 μm. Graphs show average of the number of branches, junctions and sholl analysis of microglia in control and clemastine-treted mice. **E)** Representative compressed z confocal images showing the expression of iNOS, a pro-inflammatory marker, and CD68, a phagocytic marker, in Iba1^+^ microglia in corpus callosum of control and clemastine-treated mice. Scale bar = 10 μm. Graphs show the average P2y12 and CD68 staining intensity (n= 6 mice per group). *p<0,05, **p<0,01.

Clemastine treatment did not alter the number of microglial cells or the immunoreactivity of Iba1, indicating no change in microglia population. Additionally, we examined the morphology of Iba1^+^ microglia from control and treated mice (Fig. 4C). For that, we quantified the number of branches and junctions per cell, as well as, the cellular complexity as determined by Sholl analysis. Both data demonstrated an increase in the morphology complexity and ramification in microglia from clemastine-treated mice (Fig. 4D). Since the activation state of microglia is determinant for the remyelination capacity of OPCs (Zabala et al., 2018), we wonder whether clemastine could alter microglia activation state. Clemastine induced a massive reduction in the expression of iNOs, a proinflammatory marker, as well as the expression of the phagocytic marker CD68 (Fig. 4E,F). All these data suggest that clemastine modulates microglial shape and function during development.

We further explored the expression of pro-inflammatory/anti-inflammatory genes, lipid and phagocytosis markers as well as growth factors involved in developmental myelinogenesis and adaptive myelination (Geraghty et al., 2019; Wlodarczyk et al., 2017) in FACS isolated microglia. Although we did not find changes in pro-inflammatory markers, we observed a significant upregulation of anti-inflammatory markers (Fig. 5A). In contrast, the expression of ApoE and CD36, genes involved in metabolism and phagocytosis respectively, was significantly downregulated. Notably, clemastine induced a significant reduction in *IGF* expression in microglia (Fig. 5A). Microglia CD11c^+^ has been described as a microglial subset expressed transiently in *corpus callosum* during development that play a pivotal role in myelinogenesis since it is the main source of IGF, a growth factor essential for myelinogenesis. We next analysed, using CD11c+ eYFP receporter mice, the impact of clemastine in this cell type population. Since microglia CD11c^+^ is a transient population that disappears early during development, we treated mice with clemastine from P5 to P10. Blinded analysis showed that clemastine reduced the number of microglia CD11c+ in corpus callosum in clemastine-treated mice (Fig. 5B). Altogether, these results are consistent with the hypothesis that activation of muscarinic receptors expressed in microglia with clemastinecan impair myelindevelopment.

**Figure 5.**
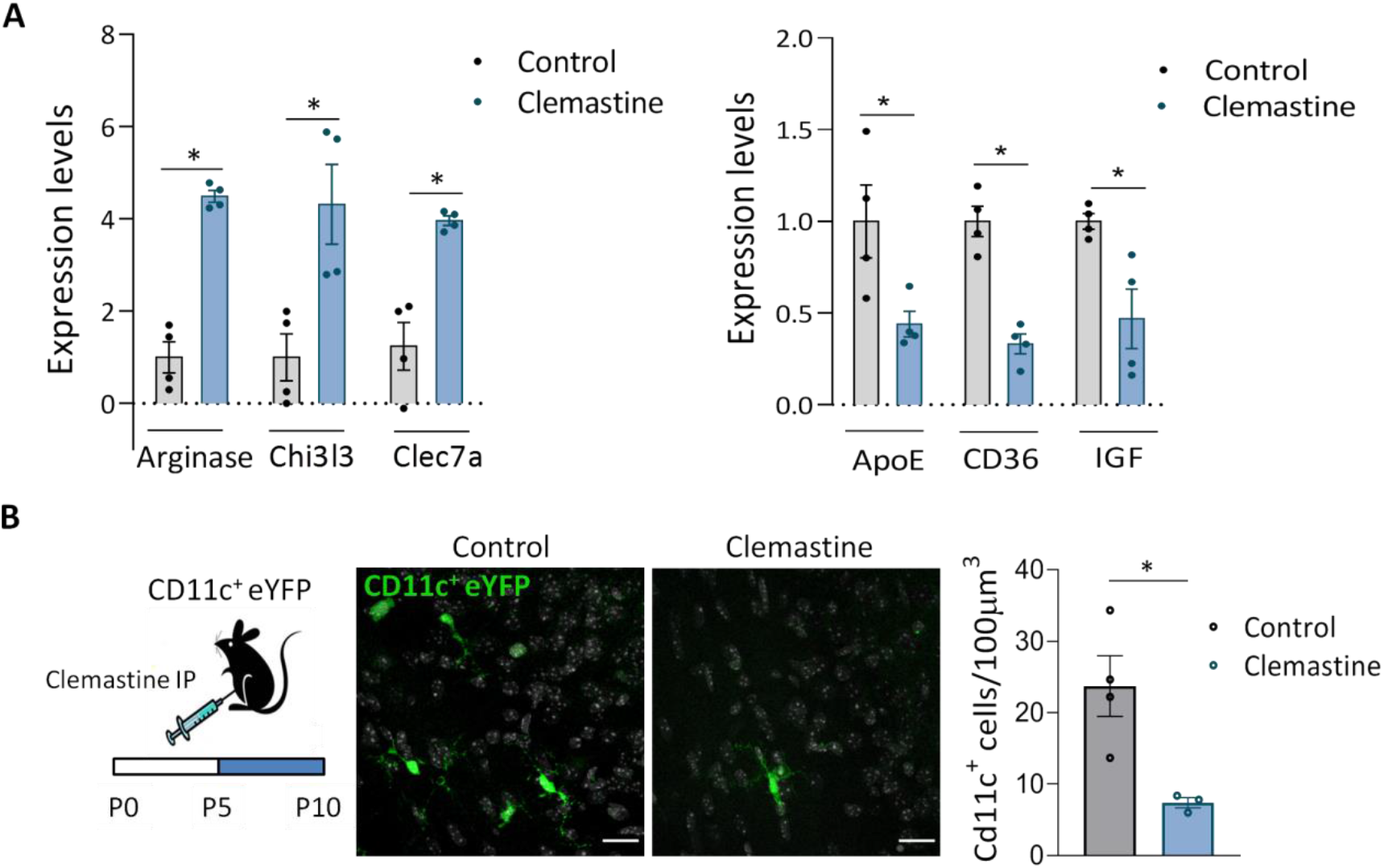
**A)** Gene expression of *Arginase*, *Chi3l3* and *clec7a*, anti-inflamatory markers, of *ApoE*, a disease associated marker, of *CD36*, a phagocytic marker and of *IGF*, a growth factor involved in myelinogenesis. The expression levels of genes are presented using fold-change values transformed to Log2 format compared to control (n= 4 mice per group). **B)** Analysis of the number of CD11c^+^ cells in control and clemastine treated CD11-eYFP mice. Representative z confocal images and histogram showing that clemastine reduced the number of CD11c^+^ cells (n= 3-4 mice per group). Scale bar = 20 μm.*p<0,05, **p<0,01

## DISCUSSION

Here we report that clemastine treatment during development promotes oligodendrocyte differentiation and alters myelination. In addition, our results also demonstrate that clemastine reduces the density of CD11c^+^ microglial cells, a transient microglia subtype present during development that it is involved in myelinogenesis. These findings strongly suggest that clemastine final outcome on myelination may depend on microglia-oligodendrocytes crosstalk.

Clemastine was identified in a high-throughput study (Mei et al., 2014) as a drug inducing oligodendrocyte progenitor differentiation *in vitro*. The drug facilitates remyelination after demyelination (Li et al., 2015; Mei et al., 2016). Thus, clemastine promotes remyelination in animal models of multiple sclerosis and after ischemic insults (Cree et al., 2018; Wang et al., 2018). Myelin renewal induced by clemastine also prevents cognitive symptoms secondary to aging and Alzheimer’s disease (Wan F et al., 2020; Chen et al., 2021). Finally, it has been described to promote myelination and to rescue behavioral phenotype in socially isolated mice (Liu et al., 2016). In contrast, no clinical benefit of clemastine was obtained in animal models of Pelizaeus Merzbacher disease (Turski et al., 2018), a dysmyelinating disorder of the central nervous system caused by impaired differentiation of oligodendrocytes during development. In accordance, we found that developmental myelination is not increased after chronic clemastine treatment, even if clemastine induced an increase in oligodendrocyte differentiation. The dose used in our study was similar to those used in other studies (Liu et al., 2016). In addition, clemastine crosses the blood brain barrier and reaches the CNS parenchyma, as determined by mass spectrometry (Turski et al., 2018).

Of note, the study of Tursky and colleagues, and ours tested the therapeutical potential of clemastine during development, whereas all the previous studies analyzed the effect of clemastine on adult mice after demyelination or altered myelination. The other important difference is that mice either do not have demyelination (in our study) or have a primary myelination disorder in Pelizaeus Merzbacher disease mice (Turski et al., 2018), whereas in the other models in which clemastine has been tested, myelination deficits were induced by toxins, inflammation or social deprivation and most of the studies are associated with chronic inflammation.

Although the beneficial effect of clemastine is extended to different insults and diseases, the underlying mechanisms are still unclear. In vitro screening proposes muscarinic M1 receptor as the target for the effect in OPCs promoting its maturation (Deshmukh et al., 2013; Mei et al., 2014). Indeed, we observe that mice treated with clemastine during development showed an increase in oligodendrocyte maturation, as revealed by the ratio NG2^+^/APC^+^. However, myelin wrapping of axons seems to be incomplete or delayed in clemastine-treated mice. In addition to oligodendrocytes, it has been reported that clemastine could interact with microglia too as it possess immune suppressive properties. Indeed, clemastine promotes neuronal protection in animal models of ALS and alleviates hypomyelination after hypoxia by modulating microglia inflammatory reaction (Apolloni et al., 2016; Xie et al., 2021).

Our study also showed that clemastine targets microglia cells during development and induced morphological changes as well as changes in the expression of inflammatory markers, a fact that could indirectly impact developmental myelination. Recent findings highlight the important role played by microglia cells in developmental myelination. Microglial cells engulf myelin sheaths during development to sculpt myelination according to axonal activity (Hughes & Appel, 2019). On the other hand, a particular CD11c^+^ microglia subset that predominates in primary myelinating areas of the developing brain is essential for myelinogenesis (Wlodarczyk et al., 2017). These cells promote myelinogenesis because they are the major source of insulin-like growth factor (IGF), a factor critical for proper myelin formation (Goebbels et al., 2010; Harrington et al., 2010). The primary signaling receptor, insulin-like growth factor-1 receptor (IGF1R), is a transmembrane tyrosine kinase receptor that binds insulin-like growth factors 1 and 2 (IGF1 and IGF2) and signals through the PI3K-AKT-mTOR, a signaling pathway whose precise regulation is critical for proper myelin formation. IGF modulates lipid metabolism (Hackett et al., 2020), a pathway essential for myelination, not for oligodendrocyte differentiation. Thus, expression of constitutively active Akt in oligodendrocytes and their progenitor cells generated no more oligodendrocytes, but dramatically more myelin, indicating that this signalling pathway could affect myelin generation without affecting OL differentiation (Flores et al., 2008). Similarly, IGF-1-overexpressing mice showed a dramatic increase in the amount of myelin per oligodendrocyte, but normal numbers of oligodendrocytes (Carson et al., 1993). More recent studies involve the Akt-mTOR signaling pathway in myelin seath growth (Fedder-Semmes & Appel, 2021). In our study we showed a significant reduction of IGF and of CD11c^+^ microglia cells in the *corpus callosum* of clemastine-treated mice. It could be possible that clemastine, under this particular circunstances, blocks the process of myelin wrapping by mature myelinating oligodendrocytes. Further experiments are necessary to confirm this hypothesis.

## Acknowledgments

This work was supported by Spanish Ministry of Education and Science (SAF2016-75292-R); Spanish Ministry of Science and Innovation (PID2019-109724RB-I00); Basque Government (PI-2016-1-0016); the University of the Basque Country (UPV/EHU); and Centro de Investigación Biomédica en Red, Enfermedades Neurodegenerativas (CIBERNED; grant CB06/05/0076). A.P. has a predoctoral fellowship from the University of the Basque Country (UPV/EHU) and A. O-A has a postdoctoral fellowship from the Basque Government. We kindly acknowledge the Microscopy Facilities of both the University of the Basque Country (UPV/EHU) and Achucarro Basque Center for Neuroscience for the technical support.

